# Dopamine influences attentional rate modulation in Macaque posterior parietal cortex

**DOI:** 10.1101/2020.05.15.097675

**Authors:** Jochem van Kempen, Christian Brandt, Claudia Distler, Mark A. Bellgrove, Alexander Thiele

## Abstract

Cognitive neuroscience has made great strides in understanding the neural substrates of attention, but our understanding of its neuropharmacology remains incomplete. Although dopamine has historically been studied in relation to frontal functioning, emerging evidence suggests important dopaminergic influences in parietal cortex. We recorded single- and multi-unit activity whilst iontophoretically administering dopaminergic agonists and antagonists while rhesus macaques performed a spatial attention task. Out of 88 units, 50 revealed activity modulation by drug administration. Dopamine inhibited firing rates according to an inverted-U shaped dose-response curve and increased gain variability. Dopamine modulated attention-related rate changes and Fano Factors in broad and narrow-spiking units, respectively. D1 receptor antagonists diminished firing rates according to a monotonic function and interacted with attention modulating gain variability in broad-spiking units. Finally, both drugs decreased the pupil light reflex. These data show that dopamine shapes neuronal responses and modulates attentional processing in parietal cortex.

## Introduction

Selective attention refers to prioritization of behaviorally relevant, over irrelevant, sensory inputs. Convergent evidence from human neuropsychological, brain imaging and non-human primate studies shows that fronto-parietal networks are crucial for selective attention (Corbetta and Shulman, 2011; Desimone and Duncan, 1995; Posner, 1990). Neuromodulation of attention-related activity in these networks occurs at least in part via glutamatergic (Dasilva et al., 2021; Herrero et al., 2013) and cholinergic inputs (Dasilva et al., 2019; Furey et al., 2008; Herrero et al., 2008; Levin and Simon, 1998; Nelson et al., 2005; Parikh et al., 2007; Sarter et al., 2005; Warburton and Rusted, 1993). Multiple lines of evidence, however, also suggest dopaminergic modulation (Bellgrove and Mattingley, 2008; Noudoost and Moore, 2011a; Soltani et al., 2013; Thiele and Bellgrove, 2018). Here we sought to understand how dopamine (DA) applied to macaque posterior parietal cortex (PPC) modulates attention-related activity.

The functional significance of DA is well established for a number of brain areas, particularly the frontal cortex (executive control) and basal ganglia (motor control). For these regions, substantial across-species similarities allowed the development of mechanistic models with clinical translational value for various disorders (e.g., Parkinson’s disease, schizophrenia or attention deficit hyperactivity disorder (ADHD)) (Arnsten et al., 2012; Thiele and Bellgrove, 2018). Species differences with respect to dopaminergic innervation do however exist for posterior cortical areas, including the PPC. Although sparse in rodents, dopaminergic innervation of parietal areas in non-human primates is comparable in strength and laminar distribution to prefrontal regions (Berger et al., 1991). Moreover, macaque PPC has high densities of DA transporter (DAT) immunoreactive axons (Lewis et al., 2001). These observations align with dense dopaminergic receptor expression in human PPC (Caspers et al., 2013) and imaging studies of clinical disorders where medications targeting DA receptors or transporters modulate parietal activity (Mehta et al., 2000). Given these data and the clinical significance of PPC function, greater understanding of dopaminergic effects in this region is warranted.

Selective attention relies heavily on PPC integrity and multiple lines of evidence suggest that DA modulates attentional processes related to parietal function. First, DA agonists reduce spatial inattention in neurological (Gorgoraptis et al., 2012) and psychiatric patients with disorders such as schizophrenia (Maruff et al., 1995) and ADHD (Bellgrove et al., 2008; Silk et al., 2014). Second, psychopharmacological studies in healthy volunteers suggest that DA antagonists modulate parameters of spatial cueing paradigms (e.g. validity effect), often associated with parietal function (Clark et al., 1989). Third, DNA variation in a polymorphism of the DA transporter gene (DAT1) is associated with individual differences in measures of spatial selective attention (Bellgrove et al., 2009, 2007; Newman et al., 2014). Fourth, non-human primate studies revealed dopaminergic contributions to working memory signals in dorsolateral prefrontal cortex (dlPFC) (Williams and Goldman-Rakic, 1995), and modulation of dopaminergic signaling in frontal eye fields (FEF) affects V4 neurons in a manner similar to attention and biases behavioral choices (Noudoost and Moore, 2011a; Soltani et al., 2013). DA thus contributes to working memory, target selection and probably also spatial attention in dlPFC and FEF (Clark and Noudoost, 2014; Noudoost and Moore, 2011a, 2011b; Williams and Goldman-Rakic, 1995). Both areas are critical nodes of fronto-parietal attention networks. In summary, while dopaminergic influences on frontal circuits are comparatively well understood, their effect on attention-related activity in PPC is yet to be established.

Here we sought to address this knowledge gap by locally infusing DA or the selective D1 receptor (D1R) antagonist SCH23390 into the PPC of two macaque monkeys during a selective attention task. We showed that single and multi-unit (SU, MU) activity is inhibited by iontophoresis of dopaminergic drugs into intraparietal sulcus (IPS) gray matter and that drug application increased trial-to-trial excitability fluctuations, termed gain variability (Goris et al., 2014). The effects of the non-selective agonist DA followed an inverted U-shaped dose-response curve, whereas the dose-response curve of the D1-selective antagonist SCH23390 followed a monotonic function. Additionally, we found cell-type specific effects on attentional modulation whereby DA affected attention-related activity and Fano Factors in broad-spiking and narrow-spiking units, respectively, whereas SCH23390 application affected attention-related gain variability changes in broad-spiking units only. Finally, both drugs reduced the pupillary light reflex.

## Results

We recorded activity from 88 single and multi-units from intraparietal sulcus (IPS) in two awake, behaving Macaque monkeys performing a selective attention task (Figure 1). Of these units, 74 (84.1%) were modulated by attention, as measured during the 500 ms before the first dimming event (see Figure 2). During recording, we used an electrode-pipette combination to iontophoretically administer dopaminergic drugs in the vicinity of the recorded cells (Thiele et al., 2006). Across the two monkeys, we recorded from 59 units whilst administering the unselective agonist DA and from 29 units during which we administered the selective D1R antagonist SCH23390. Firing rates in 36 (61%) and 14 (48.3%) units were modulated by application of DA and SCH23390, respectively. Of these drug-modulated units, 31 (52.5%) and 14 (48.3%) were also modulated by attention. Thus, approximately half the total units were modulated both by attention and drug application. These proportions are comparable to cholinergic modulation of attention induced activity in macaque V1 and FEF (Dasilva et al., 2019; Herrero et al., 2008), and glutamatergic modulation in FEF (Dasilva et al., 2021). As expected given the focal nature of micro-iontophoretic drug application (Herz et al., 1969), and in line with comparable studies (Jacob et al., 2016, 2013), there were no behavioral effects of drug application (i.e., reaction times) (Supplementary figure 1).

**Figure 1.**
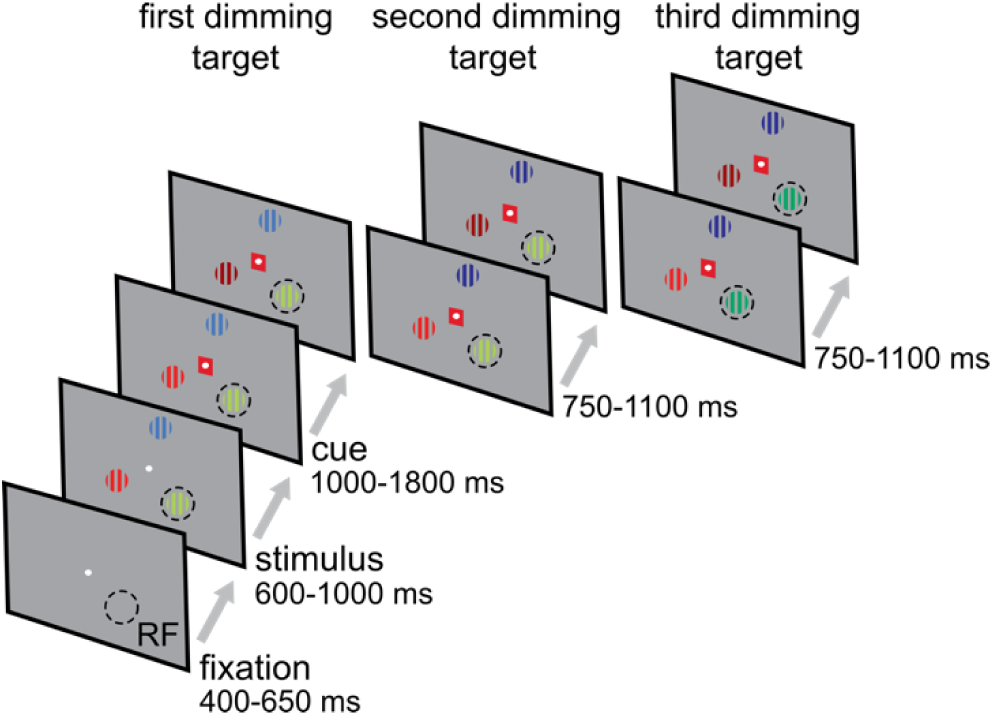
Behavioral paradigm. The monkey held a lever and fixated on a central fixation spot to initiate the trial. One of three colored gratings was presented inside the receptive field (RF) of the neurons under study. After a variable delay a cue matching one of the grating colors surrounded the fixation spot, indicating which grating was behaviorally relevant (target). In pseudorandom order the stimuli decreased in luminance (dimmed). Upon dimming of the target, the monkey had to release the lever to obtain a reward.

**Figure 2.**
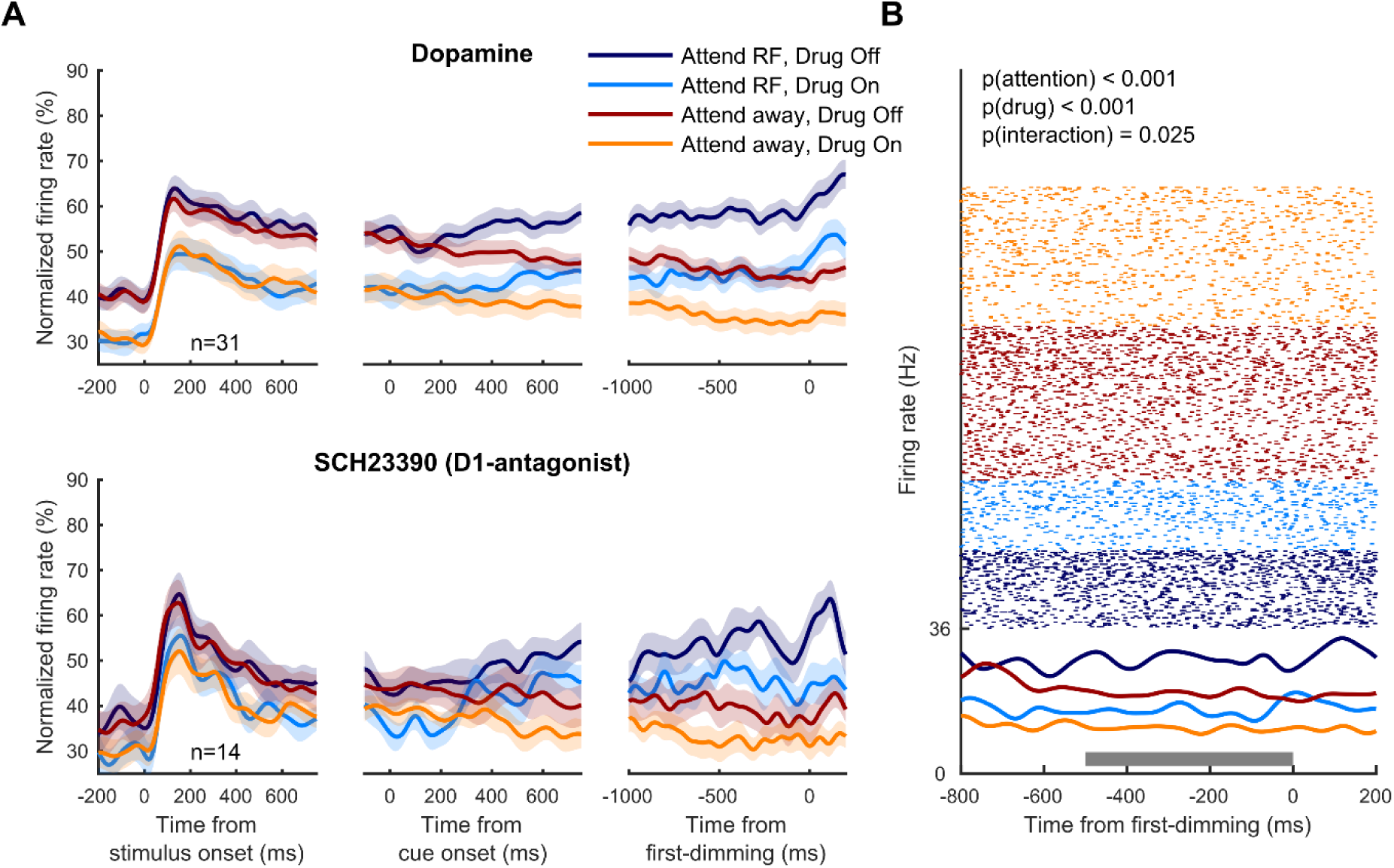
Population activity and example unit. (**A**) Population histograms for all units recorded during dopaminergic drug application selective for attention and drug application. Population activity aligned to stimulus onset (left), cue onset (middle) and the first dimming event (right), for the non-specific agonist dopamine (top) and the D1R antagonist SCH23390 (bottom). Activity is normalized for each unit by its maximum activity. Error bars denote ±1 SEM. (**B**) Activity from a representative cell recorded during dopamine application. This cell’s activity, aligned to the first dimming event, was significantly modulated by attention, drug application and showed a significant interaction between these factors. The grey bar indicates the time window used for statistical analyses. Statistics: two-factor ANVOVA.

Figure 2A illustrates the population activity (from all units) aligned to stimulus onset, cue onset and the first-dimming event, for both the no-drug and the drug conditions. For a given drug condition, neural activity between attention conditions did not differ when aligned to stimulus onset but started to diverge approximately 200 ms after cue onset, indicating which of the three gratings was behaviorally relevant on that trial, and diverged further leading up to the first dimming event. Across the population, DA strongly reduced firing rates throughout the duration of the trial, including during baseline periods as well as stimulus and cue presentation. The effects of SCH23390 were of the same sign but weaker. Control recordings (saline with matched pH) to control for pH or current related effects did not reveal any effects on firing rates (Supplementary figure 2), and thus exclude the possibility that drug effects were the result of recording or application methods. Although drug induced changes to attentional modulation of neural activity appear relatively small at the population level, a subset of neurons revealed an interaction between attention and drug application (n=9), as illustrated for an example neuron in Figure 2B, and these effects depended on the cell types affected (further delineated below). Next, we examined units that were modulated by attention and/or drug application and investigated whether activity modulation due to attention and drug application mapped onto different cell types.

Cells were classified as narrow or broad-spiking cells according to the median duration of the peak-to-trough time of the spike waveforms (Figure 3A & B). These cell types have previously been found to respond differently to dopaminergic drug application in frontal cortex (Jacob et al., 2016, 2013). Although narrow and broad-spiking cells have been argued to respectively constitute inhibitory interneurons and excitatory pyramidal cells (Mitchell et al., 2007), a more recent study found that output cells in primary motor cortex (unequivocal pyramidal cells) had a narrow action potential waveform (Vigneswaran et al., 2011), and most pyramidal cells in macaque motor cortex express the Kv3.1b potassium channel, associated with the generation of narrow spikes (Soares et al., 2017). Therefore, the narrow-broad categorization distinguishes between two different cell type categories, without mapping this classification specifically onto interneurons or pyramidal cells.

**Figure 3.**
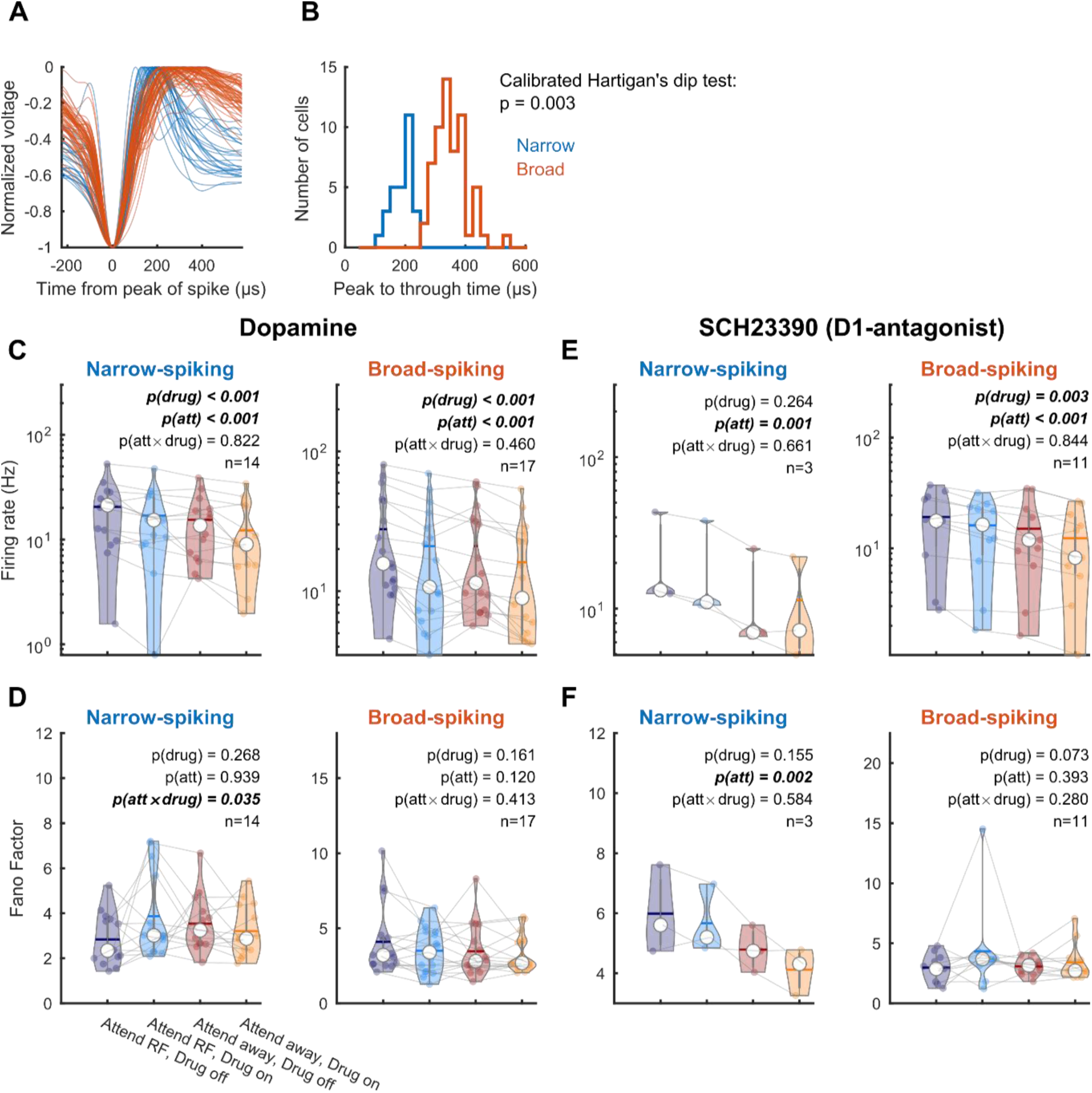
Dopaminergic modulation of firing rates across broad and narrow-spiking units. (**A**) Average spike waveforms for the population of units. (**B**) Distribution of peak-to-trough ratios. Statistics: calibrated Hartigan’s dip test (Ardid et al., 2015). (**C**) Average firing rates between attention and drug conditions for the non-specific agonist dopamine for narrow-spiking (left) and broad-spiking (right) units. (**D**) Fano factors between attention and drug conditions for the non-specific agonist dopamine. (**E**-**F)** Same conventions as (**C**-**D**) but for the D1R antagonist SCH23390. Only units that revealed a main or interaction effect for the factors drug and attention were included in this analysis. Individual markers represent the average firing rate or Fano Factor for a single unit. The white marker denotes the median and the error bars the interquartile range. Horizontal bars denote the mean. Statistics: linear mixed-effect models.

We tested whether DA application affected firing rates or rate variability, as quantified by the Fano Factors (FF) and gain variability, measured during the 500 ms preceding the first dimming, using linear mixed-effect models with categorical (effect coded) factors of drug (on/off), attention (RF/away) and unit type (narrow/broad). Confidence intervals were computed across 5000 bootstrap replicates. To control for Type I errors and to aid interpretation of model fit statistics, we additionally report the Kenward-Roger approximation for performing F tests as well as the Bayes factor (Materials & Methods). We followed these analyses with tests within each unit type, depicted in Figure 3 and Figure 4. For firing rates, we found a main effect of attention (*β* = 2.67±0.38, 95% confidence interval = [1.91, 3.45], χ^2^(1) = 29.2, P = 6.44e^-8^, P_KR_ = 8.19e^-8^, BF = 6.65e^6^) reflecting the firing rate increase when attention is directed towards the RF, and a main effect of drug (*β* = -2.31±0.38, 95% confidence interval = [-3.09 -1.55], χ^2^(1) = 31.1, P = 2.44e^-8^, P_KR_ = 3.74e^-8^, BF = 2.06e^7^), indicating that DA application reduced firing rates (Figure 3C). We did not find a main effect of unit type or any interaction. For FF, we did not find any main effects of attention, drug or unit type, but we found a trending interaction effect between drug and unit type (*β* = 0.18±0.10, 95% confidence interval = [-0.01 0.38], χ^2^(1) = 2.97, P = 0.084, P_KR_ = 0.09, BF = 1.08) and a three-way interaction between drug, attention and unit type (*β* = 0.22±0.10, 95% confidence interval = [0.03, 0.42], χ^2^(1) = 4.75, P = 0.029, P_KR_ = 0.036, BF = 3.37). This interaction reflects that when attention is directed towards the RF, DA application increases FF, whereas when attention is directed away from the RF, DA application decreases FF in narrow-spiking units (Figure 3D).

**Figure 4.**
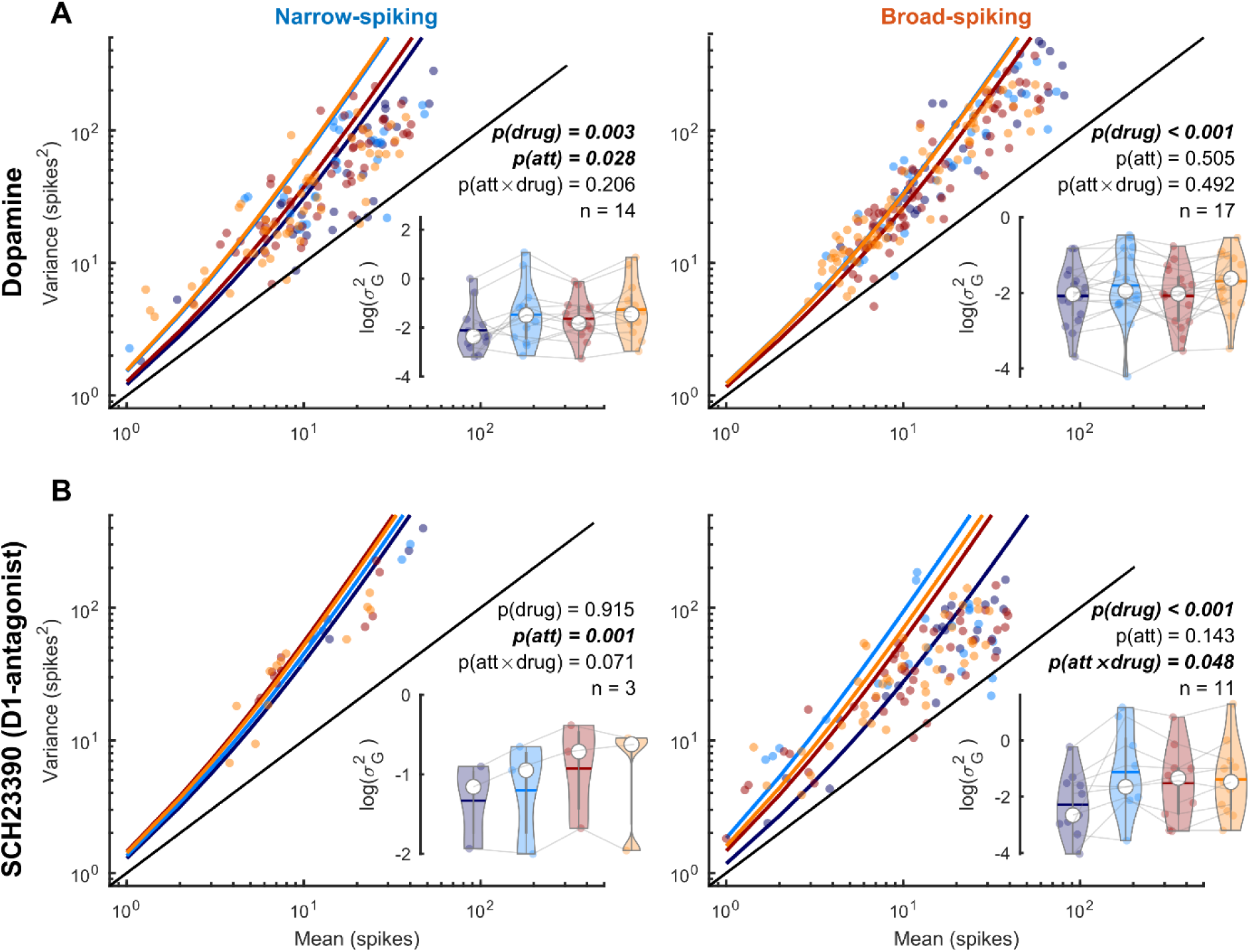
Dopaminergic modulation of gain variability across broad and narrow-spiking units. (**A**) Variance-to-mean relationship across attention and drug conditions for narrow-spiking (left) and broad-spiking (right) units for the non-specific agonist dopamine. Individual dots depict the variance and mean across trials for a single condition. Solid lines show the predicted mean-to-variance relationship given the average fitted dispersion parameter (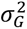). Insets show 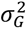 for each unit and their comparison across attention and drug conditions. Individual markers represent the gain variability for a single unit. The white marker denotes the median and the error bars the interquartile range. Horizontal bars denote the mean. (**B**) Same conventions as (**A**) but for the D1R antagonist SCH23390. Only units that revealed a main or interaction effect for the factors drug were included in this analysis. Statistics: linear mixed-effect models.

We performed the same analyses for the application of SCH23390. For firing rates, we found a main effect of attention (*β* = 3.33±0.50, 95% confidence interval = [2.33, 4.30], χ^2^(1) = 20.9, P = 4.92e^-6^, P_KR_ = 7.21e^-6^, BF = 3.22e^4^) reflecting the firing rate increase when attention is directed towards the RF, and a main effect of drug (*β* = -1.29±0.50, 95% confidence interval = [-2.3, -0.29], χ^2^(1) = 8.47, P = 0.004, P_KR_ = 0.005, BF = 13.3), indicating that DA application reduced firing rates (Figure 3E). We additionally found an interaction between attention and unit type (*β* = 1.35±0.50, 95% confidence interval = [0.37, 2.33], χ^2^(1) = 6.72, P = 0.01, P_KR_ = 0.014, BF = 4.9), indicating that narrow-spiking units increased their firing rates more when attention was directed towards the RF. We did not find any effect of drug application or attention for FF, but we found a trending main effect of unit type (*β* = 0.85±0.42, 95% confidence interval = [-0.002, 1.69], χ^2^(1) = 3.49, P = 0.06, P_KR_ = 0.09, BF = 0.19). However, the lack of clear significant effects in conjunction with the low number of narrow-spiking units for this sample raise doubts about their robustness (Figure 3F).

We next investigated the effects of drug application and attention on gain variability (Goris et al., 2014). Neural activity often displays super-Poisson variability (larger variance than the mean), resulting from trial-to-trial changes in excitability, that can be modeled by fitting a negative binomial distribution to the spike rate histogram. This distribution is characterized by a dispersion parameter that captures this additional variability and has been proposed to reflect stimulus-independent modulatory influences on excitability (Goris et al., 2014). Whereas FF is a measure of variability that is accurate when the variance is proportional to the mean, gain variability captures the nonlinear variance-to-mean relationship (Thiele et al., 2016). During DA application we found a trending main effect of attention (*β* = -0.1±0.041, 95% confidence interval = [-0.18, -0.02], χ^2^(1) = 3.26, P = 0.07, P_KR_ = 0.07, BF = 0.6) and a main effect of drug application (*β* = 0.20±0.041, 95% confidence interval = [0.12, 0.28], χ^2^(1) = 18.5, P = 1.72e^-5^, P_KR_ = 2.33e^-5^, BF = 1.38e^4^) on gain variability. This indicates increased variability during drug application and decreased variability when attention was directed towards the RF. We furthermore found a trending interaction between attention and unit type (*β* = -0.07±0.041, 95% confidence interval = [-0.15, 0.01], χ^2^(1) = 2.72, P = 0.099, P_KR_ = 0.11, BF = 0.65), revealing a decrease in gain variability in narrow-spiking units when attention was directed towards the RF (Figure 4A). For SCH23390, we found a trending main effect of attention (*β* = -0.14±0.081, 95% confidence interval = [-0.3, 0.02], χ^2^(1) = 3.52, P = 0.061, P_KR_ = 0.065, BF = 1.08) and a main effect of drug application (*β* = 0.16±0.081, 95% confidence interval = [0.0004, 0.32], χ^2^(1) = 9.04, P = 0.003, P_KR_ = 0.004, BF = 37), indicating increased gain variability with drug application and decreased variability when attention was directed towards the RF. In addition, there was as a trending interaction effect between drug application and unit type (*β* = 0.16±0.081, 95% confidence interval = [-0.31, 0.001], χ^2^(1) = 3.56, P = 0.059, P_KR_ = 0.08, BF = 1.33), indicating a relatively larger difference in gain variability in broad compared to narrow-spiking units. The model fits within each unit type revealed a significant main effect of drug application (*β* = 0.31±0.088, p = 0.0009) and an interaction between drug application and attention (*β* = 0.18±0.088, p = 0.048) for broad-spiking units. For narrow-spiking units we found a main effect of attention (*β* = -0.14±0.03, p = 0.001) and a trending interaction effect between drug application and attention (*β* = 0.06 ±0.03, p = 0.071) (Figure 4B).

To investigate whether DA affected attention-specific activity, we tested if attention AUROC values were modulated by drug application. Attention AUROC values indicate how well an ideal observer can distinguish between neural activity during attend RF or attend away trials. A value of 0.5 indicates that the distributions are indistinguishable, whereas values of 0 or 1 indicate perfectly distinguishable distributions. Drug application reduced AUROC values for broad-spiking cells, whereas narrow-spiking cells were unaffected (Figure 5A) [two-sided Wilcoxon signed-rank test; narrow-spiking: Δ-AUROC -0.002±0.01, p=0.952, Cohen’s d=0.030; broad-spiking: Δ-AUROC -0.034±0.006, p=0.009, Cohen’s d=-0.70]. Corrected AUROC values (1-AUROC if the AUROC value was smaller than 0.5 without drug application, Materials & Methods) revealed a trending relationship [two-sided Wilcoxon signed-rank test; broad-spiking: Δ-AUROC -0.02±0.01, p=0.08, Cohen’s d=-0.38]. SCH23390 application did not modulate AUROC values for either cell type (Figure 5B). DA thus had a cell-type specific effect on attentional rate modulation, but this was only trending, once corrected values of AUROCs were used.

**Figure 5.**
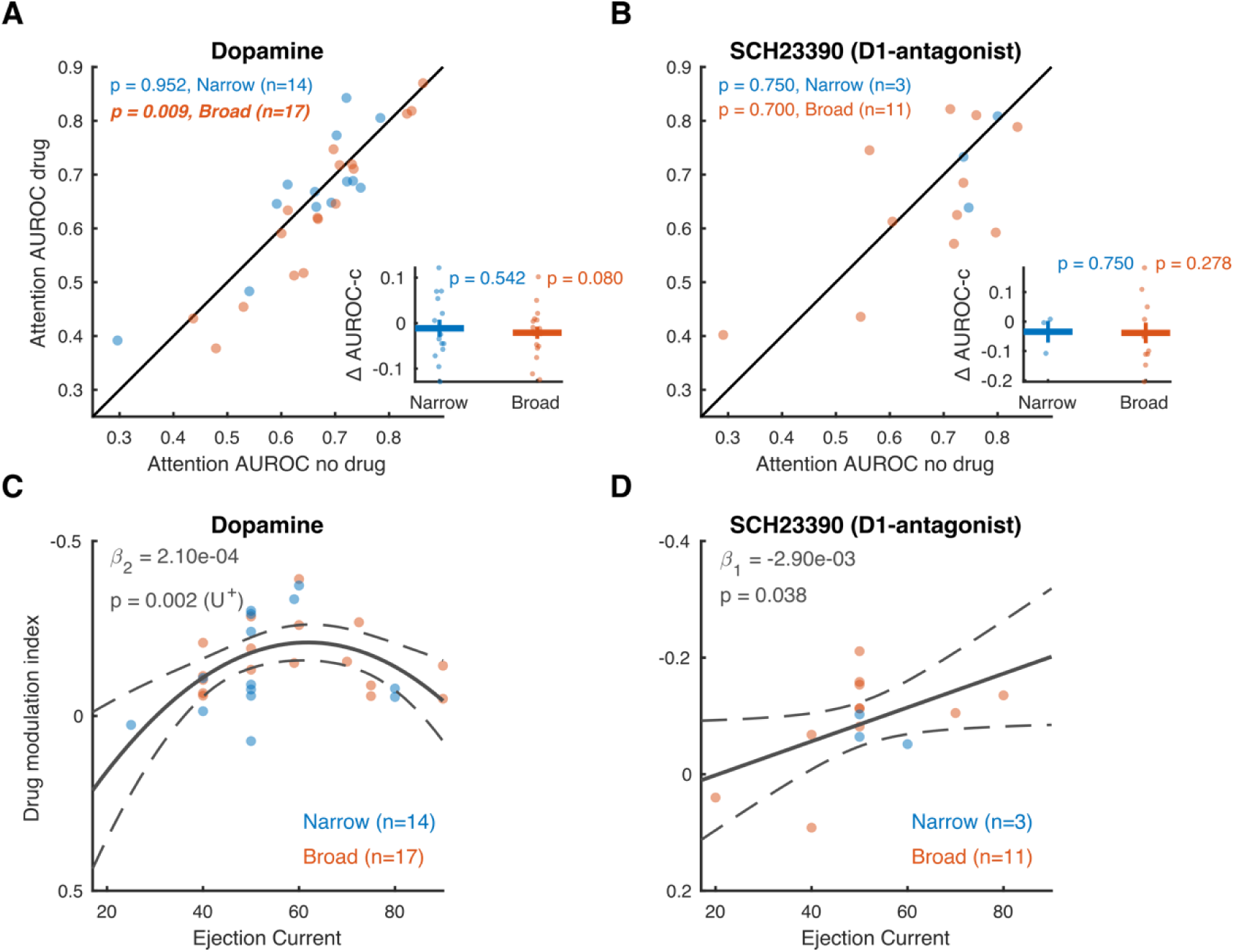
Dopaminergic modulation of AUROC values and dose-response curves. (**A**-**B**) Area under the receiver operating characteristic (AUROC) curve between no drug and drug conditions for the non-specific agonist dopamine (**A**) and the D1R antagonist SCH23390 (**B**). The insets depict the difference (drug-no drug) of the corrected AUROC values (Materials & Methods). Only cells that revealed a main or interaction effect for the factors of drug and attention were included in this analysis. Statistics: Wilcoxon signed rank tests (FDR corrected). Statistics deemed significant after multiple comparison correction are displayed in italic and boldface fonts. (**C**-**D**) Drug modulation index plotted against ejection current for the non-specific agonist dopamine (**C**) and the D1R antagonist SCH23390 (**D**). Note the reversed y-axis. Solid and dotted lines represent significant model fits (applied to all cells simultaneously) and their 95% confidence intervals, respectively. A monotonic relationship is shown if a first-order fit was better than a constant fit, and a non-monotonic relationship is shown if a second-order fit was better than a linear fit. U^+^ indicates a significant U-shaped relationship. Only cells that revealed a main or interaction effect for the factor drug were included in this analysis. Statistics: linear mixed-effects model analysis.

We applied dopaminergic drugs with a variety of iontophoretic ejection currents (20-90 nA). Since DA has previously been shown to modulate neural activity according to an inverted U-shaped dose-response curve (Vijayraghavan et al., 2007), with maximal modulation at intermediate DA levels, we tested whether the ejection current was predictive of the firing rate modulation associated with drug application, estimated by a drug modulation index (MI_drug_, Materials & Methods). Specifically, we used sequential linear mixed-effect model analyses and likelihood ratio tests to test for linear and quadratic trends. U-shaped trends were verified using the two-lines approach (Materials & Methods). DA displayed a non-monotonic relationship with MI_drug_ (χ^2^(1) = 9.89, p = 0.002) and revealed an inverted U-shaped curve (p < 0.05) in which intermediate ejection currents elicited the most negative MI_drug_, i.e. the largest inhibition of activity (Figure 5C). For SCH23390, on the other hand, we found a monotonic dose-response relationship (χ^2^(1) = 4.31, p = 0.038), with more inhibition of firing rates with higher drug ejection currents (Figure 5D). Neither of these dose-response relationships were dependent on unit sub-selection based on their attention or drug selectivity (Supplementary figure 3).

To investigate whether drug dosage was also predictive of attentional rate modulation, we performed the same analysis on the difference score (drug – no drug) of attention AUROC values. Neither DA (χ^2^(1) = 0.95, p = 0.330), nor SCH23390 (χ^2^(1) = 0.33, p = 0.568) dosage were predictive of attention AUROC, regardless of unit sub-selection (Supplementary figure 4).

Interestingly, we found that the application of both DA and SCH23390 influenced pupil diameter. We conducted a sliding-window Wilcoxon signed rank test analysis for each 200 ms window, in 10 ms increments, comparing baseline-normalized pupil diameter on drug compared to no-drug trials (Figure 6A). This analysis revealed a significant difference in pupil diameter that started after stimulus onset and lasted until after cue onset. Specifically, we found a small but significant modulation of the pupillary light reflex (Figure 6). The magnitude of the constriction of the pupil upon stimulus onset was reduced during dopaminergic drug application compared to control trials [two-sided Wilcoxon signed-rank test; DA: Δ-pupil 0.10±0.02, p<0.001, Cohen’s d=1.09; SCH23390: Δ-pupil 0.10±0.03, p=0.004, Cohen’s d=0.79], but neither drug influenced pupil diameter during any other time window (Figure 6B-E). Another sliding window analysis using a two factor (drug by attention) repeated measures ANOVA revealed no effect of attention (main or interaction) on pupil diameter (data not shown). Thus, locally applied dopaminergic drugs in parietal cortex modulated the pupillary light reflex upon stimulus onset.

**Figure 6.**
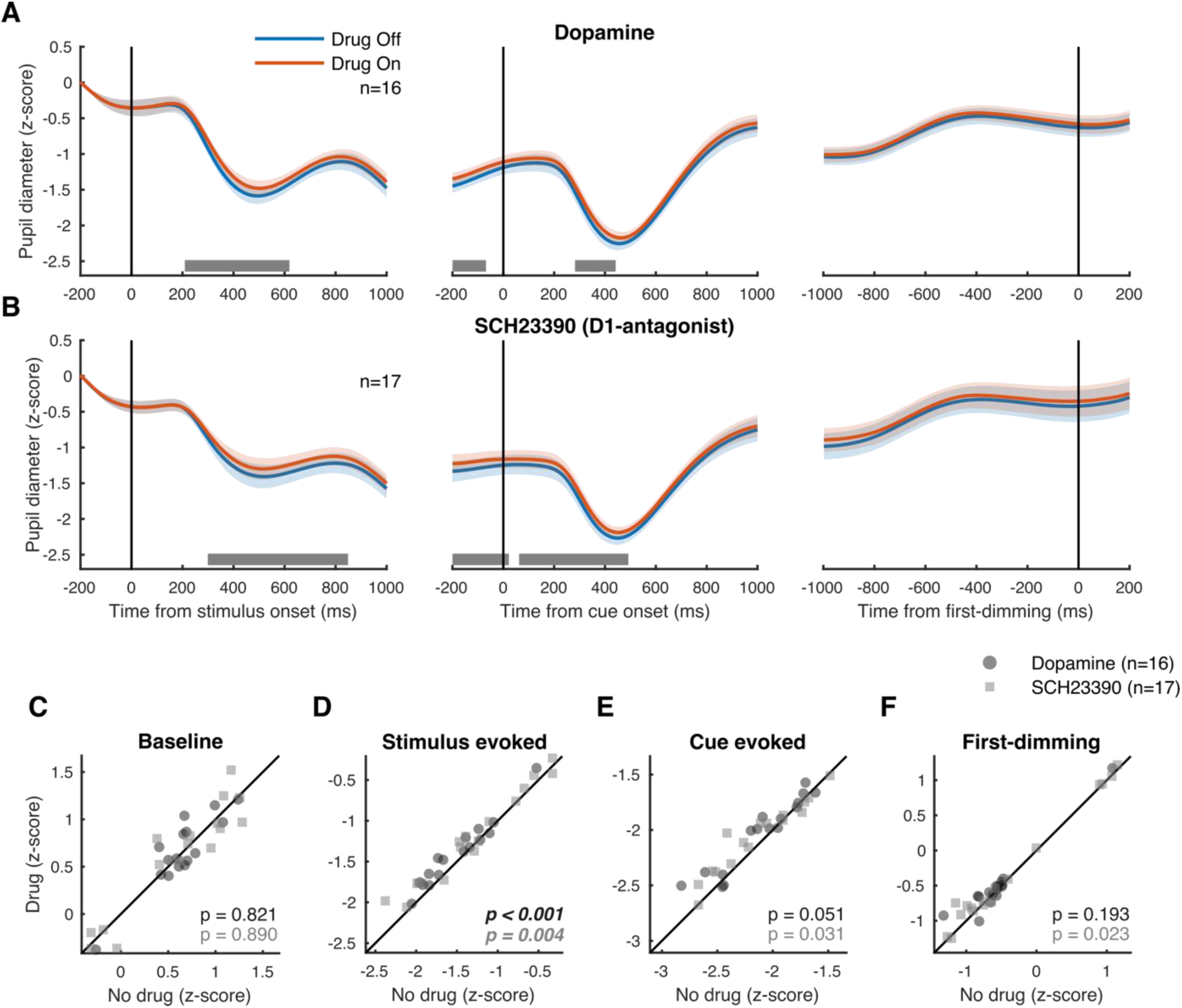
Modulation of pupil diameter by dopamine in Parietal cortex. (**A**) Pupil diameter during sessions where dopamine was administered aligned to stimulus onset (left), cue onset (middle) and the first dimming event (right). The grey bar indicates the times where drug application brought about a significant difference in pupil diameter. (**B**) As (A) but for sessions where D1R antagonist SCH23390 was applied. (**C**-**F**) Average pupil diameter during pre-stimulus baseline period (**C**), after stimulus onset (**D**), after cue onset (**E**), and before the first dimming event (**F**). Shaded regions denote ±1 SEM. Statistics: Wilcoxon signed rank test (FDR corrected). Statistics deemed significant after multiple comparison correction are displayed in italic and boldface fonts.

## Discussion

We tested the effects of dopaminergic drugs on PPC activity during spatial selective attention. The non-specific agonist DA inhibited activity according to an inverted U-shaped dose-response curve, whereas the D1R antagonist SCH23390 decreased firing rates for broad-spiking units following a monotonic dose-response curve. We found interaction effects between DA application and attention for Fano Factors in narrow-spiking units, as well as application of SCH23390 and gain variability in broad-spiking units. We further found preliminary evidence that DA reduces attention-related firing rate modulations in broad-spiking units. Finally, we found that local drug application in parietal cortex decreased the pupillary light reflex. This is the first study (to the best of our knowledge) revealing the role of dopaminergic modulation on task-related activity in the parietal cortex of the rhesus macaque.

### General and cell-type specific dopaminergic modulation in parietal cortex

We distinguished between broad and narrow-spiking units. Even though, as discussed above, this classification does not reflect a one-to-one mapping onto interneurons and pyramidal cells, this categorization may explain some of our results (Jacob et al., 2016, 2013). DA has a well-established role in modulating prefrontal signaling, supporting cognitive functions such as working memory and attention (Clark and Noudoost, 2014; Noudoost and Moore, 2011b; Ott and Nieder, 2019; Thiele and Bellgrove, 2018; Vijayraghavan et al., 2007; Watanabe et al., 1997; Williams and Goldman-Rakic, 1995). D1R and D2R are expressed broadly throughout the cortex and fulfil complementary roles in prefrontal cognitive control (Ott and Nieder, 2019). Although D2Rs have been implicated in rule coding (Ott et al., 2014), modulation of working memory is mostly associated with D1R stimulation or blockade (Sawaguchi et al., 1990; Sawaguchi and Goldman-Rakic, 1991, 1994; Williams and Goldman-Rakic, 1995). Moreover, while manipulation of either receptor subtype in FEF can modulate behavioral choices (Soltani et al., 2013), only D1R blockade in FEF elicits activity resembling attentional effects in extrastriate visual areas (Noudoost and Moore, 2011a).

Interestingly, D1R expression is higher in FEF pyramidal cells compared to interneurons (Mueller et al., 2019, 2018). Here, the effects of dopaminergic drugs were greater for broad-, rather than narrow-spiking units. Although it is unknown whether DA receptor expression differs across cell types in PPC, if expression is similar to the FEF, modulation of parietal attentional signals might rely on higher expression of D1R compared to D2R in broad-spiking putative pyramidal cells.

It is remarkable that the majority of the recorded neurons were inhibited by DA and SCH23390 application, as previous studies (in prefrontal cortex) found mixed responses to unselective DA (Jacob et al., 2013) or D1R stimulation (Vijayraghavan et al., 2007; Williams and Goldman-Rakic, 1995). As control recordings using saline did not result in any systematic effects (Supplementary figure 2), these effects were not due to our recording/iontophoresis methods.

The effects found may alternatively be explained by drug dosages. Although Jacob et al. (2013) found that the proportion of inhibited and excited cells did not differ across a variety of ejection currents (25-100 nA), activity increases have been found for low, and decreases for high D1R agonist and antagonist dosages (Vijayraghavan et al., 2007; Williams and Goldman-Rakic, 1995). Indeed, while our sample size using lower dosages was small, lower ejection currents predicted positive and less negative modulation. At the dosages used in this study, DA could have mostly inhibitory effects. Vijayraghavan et al. (2007) found that low doses (10-20 nA) of D1R agonists reduced overall firing rates, but increased spatial specificity of prefrontal neurons, whereas high dosages (20-100 nA) further reduced activity and abolished spatially selective information. Given that our study was unrelated to spatial specificity (i.e. saccade field tuning), we were unable to assess this particular feature, but dopaminergic influences may still enhance spatial tuning of PPC despite an overall reduction in activity.

Another factor that could explain the low number of DA-excited units is the short block duration used in our task. Cells excited by DA respond more slowly to drug application than inhibited cells, with an average modulation up-ramp time constant of 221.9 s (Jacob et al., 2013). In our task, with a median trial duration of approximately 8 s, a block (36 trials) lasted approximately 288 s. DA-excited neurons could have only started to show modulation towards the end of the block, resulting in a population of largely inhibited units.

In sum, dopaminergic effects on (task-related) activity are complex (Seamans and Yang, 2004) and depend on various factors not controlled for in this study, such as endogenous levels of DA. Within prefrontal cortex, coding can be enhanced by D1R agonists, and diminished by antagonists (Ott et al., 2014; Vijayraghavan et al., 2007), or vice-versa (Noudoost and Moore, 2011a; Williams and Goldman-Rakic, 1995). Indeed, dopaminergic effects show regional variability across different brain areas, even within PFC (Arnsten et al., 2012). Thus, the mechanisms discussed above might not apply to PPC. Finally, as SCH23390 also has high affinity agonistic properties for 5-HT_2c_ (serotonin) receptors (Millan et al., 2001), some of our effects might be unrelated to dopaminergic functioning. Although the effects on attention were modest and our sample size was relatively small, these results encourage future studies with larger sample sizes and a more detailed distinction between cell types to explore cell-type and receptor-subtype specific (dose-dependent) effects of DA in parietal cortex during task performance.

### Dopaminergic dose-response curve

DA receptor stimulation follows an inverted-U shaped dose-response curve whereby too little or too much stimulation leads to suboptimal behavioral performance (Arnsten et al., 1994; Zahrt et al., 1997) or neural coding (Vijayraghavan et al., 2007). Whereas optimal levels of DA receptor stimulation can stabilize and tune neural activity, suboptimal levels decrease neural coding and behavioral performance.

Here we found an inverted-U shaped dose-response curve for DA, and a monotonic function for SCH23390. Rather than predicting neural coding for attention, however, ejection currents were merely predictive of drug modulation indices, without any relationship to attention AUROC values. However, these results should be interpreted with caution. First, our sample size, especially for SCH23390, might have been too small to reliably determine the shape of the dose-response curve. Second, since lower and higher ejection currents were not used as often as intermediate currents, it is possible we did not have sufficient data to constrain the function fit at the extremes. Finally, we applied different ejection currents across rather than within cells. Based on these data, it is therefore not possible to conclusively state that individual cells in parietal cortex respond according to a U-shaped dose response curve. It is furthermore important to note that the dopaminergic effects might partly be driven by receptor subtypes (e.g. D2R) not usually associated with modulation of delay period activity. Despite these notes of caution, we believe this study provides evidence for a role of DA in parietal cortex during cognitive tasks and presents opportunities for future research to elucidate the exact underlying mechanisms.

### Dopaminergic modulation of the pupil light reflex

The pupil light reflex (PLR) transiently constricts the pupil after exposure to increases in illumination or presentation of bright stimuli (Loewenfeld, 1993; McDougal and Gamlin, 2014). Recent studies have shown that covert attention can modulate this behavioral reflex (Binda and Murray, 2015a, 2015b; Naber et al., 2013). Subthreshold FEF microstimulation respectively enhances or reduces the PLR when a light stimulus is presented inside or outside the saccade field (Ebitz and Moore, 2017). The PLR thus depends both on luminance changes and the location of spatial attention. We found that dopaminergic drug application in parietal cortex reduced the PLR. These results are in agreement with the electrophysiological results, as drug administration also reduced attentional rate modulation. Two (non-exclusive) mechanisms have been proposed by which FEF can modulate the PLR (Binda and Gamlin, 2017); by direct or indirect projections to the olivary pretectal nucleus, or via indirect projections to constrictor neurons in the Edinger-Westphal nucleus. For the latter, these projections are hypothesized to pass through extrastriate visual cortex and/or the superior colliculus (SC). Subthreshold microstimulation of the intermediate (SCi), but not superficial (SCs), layers of the SC elicits a short latency pupillary dilation (Joshi et al., 2016; Wang et al., 2012). Whereas the SCs receive input from early visual areas, including the retina, the SCi receives input from higher-order association cortices. Along with preparing and executing eye movements, the SCi is involved in directing covert attention (Ignashchenkova et al., 2004; Kustov and Lee Robinson, 1996; Lovejoy and Krauzlis, 2010; Muller et al., 2005), and provides an essential contribution to the selection of stimuli amongst competing distractors (McPeek and Keller, 2004, 2002; reviewed in Mysore and Knudsen, 2011). Moreover, the SC receives dense projections from parietal cortex (Becker, 1989; Kuypers and Lawrence, 1967), and has been hypothesized to play an important role in pupil diameter modulation (Wang and Munoz, 2015). It is currently unclear whether dopaminergic modulation of frontal (or parietal) cortex modulates SC activity, but this pathway seems a strong candidate for the modulation of the PLR (Wang and Munoz, 2015) that we encountered in this study through DA application. Here, dopaminergic drug application reduced parietal activity and brought about a gain modulation (reduction) of a brainstem-mediated reflex to fixed visual input. Although covert attention was not directed at any specific stimulus at the time of stimulus onset, the modulation of the PLR observed here is consistent with previously reported effects of covert attention and FEF microstimulation on the PLR. Speculatively, this modulation could affect the bottom-up attentional capture by the stimulus, but further studies are required to test this hypothesis.

## Conclusion

DA is an important modulator of high-level cognitive functions, both in the healthy and ageing brain as well as for various clinical disorders (Arnsten et al., 2012; Robbins and Arnsten, 2009; Thiele and Bellgrove, 2018). Although dopaminergic effects within PFC have been elucidated in some detail, the effects of DA in other brain areas such as parietal cortex, despite its well-established role in cognition and cognitive dysfunction, has largely been overlooked. This study is the first to show dopaminergic modulation of parietal activity in general, and activity specific to spatial attention in the non-human primate. Our work encourages future studies of dopaminergic involvement in parietal cortex, thereby gaining a broader understanding of neuromodulation in different networks for cognition.

## Materials & Methods

### Procedures

All animal procedures were approved by the Newcastle University Animal Welfare Ethical Review Board and performed in accordance with the European Communities Council Directive RL 2010/63/EC, the National Institute of Health’s Guidelines for the Care and Use of Animals for Experimental Procedures, and the UK Animals Scientific Procedures Act. Animals were motivated to engage in the task through fluid control at levels that do not affect animal physiology and have minimal impact on psychological wellbeing (Gray et al., 2016).

### Surgical preparation

The monkeys were implanted with a head post and recording chambers over the lateral intraparietal sulcus under sterile conditions and under general anesthesia. Surgery and postoperative care conditions have been described in detail previously (Thiele et al., 2006).

### Behavioral paradigms

Stimulus presentation and behavioral control was regulated by Remote Cortex 5.95 (Laboratory of Neuropsychology, National Institute for Mental Health, Bethesda, MD). Stimuli were presented on a cathode ray tube (CRT) monitor at 120 Hz, 1280 × 1024 pixels, at a distance of 54 cm.

The location of the saccade field (SF) was mapped using a visually- or memory-guided saccade task. Here, monkeys fixated centrally for 400 ms after which a saccade target was presented in one of nine possible locations (8-10° from fixation, distributed equidistantly). After a random delay (800-1400 ms, uniformly distributed) the fixation point was extinguished, which indicated to the monkey to perform a saccade towards the target. In the memory-guided version of the task (used only for saline-control recordings), the visual target was briefly presented in one of four locations. After extinguishing the target, its location needed to be remembered until a saccade was made towards the remembered location (after extinguishing of the fixation point). Online analysis of visual, sustained and saccade related activity determined an approximate SF location which guided our subsequent receptive field (RF) mapping. The location and size of RFs were measured as described previously (Gieselmann and Thiele, 2008), using a reverse correlation method. Briefly, during fixation, a series of black squares (1-3° size, 100% contrast) were presented for 100 ms at pseudorandom locations on a 9 × 12 grid (5-25 repetitions for each location) on a bright background. RF eccentricity ranged from 2.5° to 17° and were largely confined to the contralateral visual field.

The main task and stimuli have been described previously (Ferro et al., 2021; Thiele et al., 2016; van Kempen et al., 2021). In brief, the monkey initiated a trial by holding a lever and fixating a white fixation spot (0.1°) displayed on a grey background (1.41 cd/m^2^). After 425/674 ms [monkey 1/monkey 2] three colored square wave gratings (2° - 6°, dependent on RF size and distance from fixation) appeared equidistant from the fixation spot, one of which was centered on the RF of the recorded neuron. Red, green and blue gratings (see Table 1 for color values) were presented with an orientation at a random angle to the vertical meridian (the same orientation for the three gratings in any given session). The locations of the colors, as well as the orientation, were pseudorandomly assigned between recording sessions and held constant for a given recording session. Gratings moved perpendicular to the orientation, whereby the direction of motion was pseudorandomly assigned for every trial. After a random delay (570-830/620-940 ms [monkey 1/monkey 2], uniformly distributed in 1 ms steps) a central cue appeared that matched the color of the grating that would be relevant on the current trial. After 980-1780/1160-1780 ms [monkey 1/monkey 2] (uniformly distributed in 1 ms steps), one pseudorandomly selected grating changed luminance (dimmed). If the cued grating dimmed, the monkey had to release the lever to obtain a reward. If a non-cued grating dimmed, the monkey had to ignore this and wait for the cued grating to dim. This could happen when the second or third grating changed luminance (each after 750-1130/800-1130 ms [monkey 1/monkey 2], uniformly distributed in 1 ms steps). Drugs were administered in blocks of 36 trials. The first block was always a control block. Thereafter, drug blocks and recovery blocks were alternated until the animal stopped working (number of block reversals, median ± interquartile range = 12 ± 6).

**Table 1.** Color values used for the 3 colored gratings across recording sessions and subjects, indicated as [RGB] – luminance (cd/m^2^). a = Undimmed values, b = dimmed values.

|  | *Red* | *Green* | *Blue* |
| --- | --- | --- | --- |
| ***Monkey 1*** | a. [255 0 0] - 14.5 | a. [0 128 0] – 9.1 | a. [60 60 255] - 11.5 |
| ***Early recordings (n=29)*** | b. [100 0 0] - 1.4 | b. [0 70 0] – 1.9 | b. [10 10 140] – 2.2 |
| ***Monkey 2*** | a. [220 0 0] – 12.8 | a. [0 135 0] – 12.9 | a. [60 60 255] – 12.2 |
| ***Early recordings (n=5)*** | b. [180 0 0] – 7.7 | b. [0 110 0] – 7.3 | b. [35 35 220] – 7.4 |
| ***Monkey 1/2 (n=12/8)*** | a. [220 0 0] – 12.8 | a. [0 135 0] – 12.9 | a. [60 60 255] – 12.2 |
| ***Late recordings*** | b. [140 0 0] – 4.2 | b. [0 90 0] – 4.6 | b. [30 30 180] – 4.6 |

### Identification of recording sites

The location of the IPS was initially guided by means of postoperative structural magnetic resonance imaging (MRI), displaying the recording chamber. During each recording, neuronal response properties were determined using SF and RF mapping tasks. During the SF mapping task, we targeted cells that showed spatially selective persistent activity and preparatory activity before the execution of a saccadic eye movement.

### Electrode-pipette manufacturing

We recorded from the lateral (and in a few occasions medial) bank of the IPS using custom-made electrode-pipettes that allowed for simultaneous iontophoretic drug application and extracellular recording of spiking activity (Thiele et al., 2006). The location of the recording sites in one of the monkeys was verified in histological sections stained for cyto- and myeloarchitecture (Distler and Hoffmann, 2001).

The manufacture of the electrodes was similar to the procedures described by Thiele et al., (2006), with minor changes to the design in order to reach areas deeper into the IPS, such as the ventral part of the lateral intraparietal area (LIPv). We sharpened tungsten wires (125 µm diameter, 75 mm length, Advent Research Materials Ltd., UK) by electrolytic etching of the tip (10-12 mm) in a solution of NaNO_2_ (172.5 g), KOH (85 g) and distilled water (375 ml). We used borosilicate glass capillaries with three barrels (custom ordered, Hilgenberg GmBH, www.hilgenberg-gmbh.de), with the same dimensions as those described previously (Thiele et al., 2006). The sharpened tungsten wire was placed in the central capillary and secured in place by bending the non-sharpened end (approximately 10 mm) of the wire over the end of the barrel. After marking the location of the tip of the tungsten wire, shrink tubing was placed around the top and bottom of the glass. The glass was pulled around the tungsten wire using a PE-21 Narishige microelectrode puller with a heating coil made from Kanthal wire (1 mm diameter, 13 loops, inner loop diameter 3 mm) and the main (sub) magnet set to 30 (0) and the heater at 100. The electrode-pipette was placed such that the tip of the tungsten wire protruded 11 mm from the bottom of the heating coil. After pulling, we filled the central barrel (with the tungsten electrode inside) with superglue using a syringe and fine flexible injection cannula (MicroFil 28 AWG, MF28G67-5, World Precision Instruments, Ltd.). We found that if we did not fill (most of) the central barrel with superglue after pulling, the recorded signal was often very noisy, possibly due to small movements of the animal (such as drinking), which caused the free tungsten wire to resonate inside the glass. Using a micro grinder (Narishige EG-400), we removed excess glass, sharpened the tip of the electrode and opened the flanking barrels of the pipette. This pulling procedure resulted in a pulled electrode part of approximately 2.5 cm length, with gradually increasing diameter, from ∼10 μm to ∼200 μm, over the first 12 mm of the electrode-pipette.

### Electrode-pipette filling and iontophoresis

Electrode-pipettes were back-filled with the same drug in both pipettes using a syringe, filter units (Millex® GV, 22 μm pore diameter, Millipore Corporation) and fine flexible injection cannula (MicroFil 34 AWG, MF34G-5, World Precision Instruments, Ltd.). The pipettes were connected to the iontophoresis unit (Neurophore-BH- 2, Medical systems USA) with tungsten wires (125 μm diameter) inserted into the flanking barrels. Because of the exploratory nature of these recordings (it is unknown whether DA influences parietal neurons during spatial attention tasks and what modulation can be expected with different amounts of drug applied), we used a variety of iontophoretic ejection currents (20 - 90 nA). The choice of current was not based on the characteristics of individual cells (e.g. their responsiveness to the drug). A fixed ejection current of 50 nA was used for the saline-control recordings. The details regarding concentration and pH of the drugs were: DA (0.1M in water for injections, pH 4-5), SCH23390 (0.005-0.1M in water for injections, pH 4-5) and saline with citrate/hydrochloric acid buffer solution (pH 4). We excluded the first two trials after a block change to allow the drugs to wash in/out and avoid sudden rate changes.

### Data acquisition

Stimulus presentation, behavioral control and drug administration was regulated by Remote Cortex 5.95 (Laboratory of Neuropsychology, National Institute for Mental Health, Bethesda, MD). Raw data were collected using Remote Cortex 5.95 (1-kHz sampling rate) and by Cheetah data acquisition (32.7-kHz sampling rate, 24-bit sampling resolution) interlinked with Remote Cortex 5.95. Data were replayed offline, sampled with 16-bit resolution and band-pass filtered (0.6-9 kHz). Spikes were sorted manually using SpikeSort3D (Neuralynx). Eye position and pupil diameter were recorded using a ViewPoint eyetracker (Arrington research) at 220 Hz. Pupil diameter was recorded in 33 out of 47 recording sessions.

### Pupillometry

Pupil diameter was low pass filtered (10 Hz) using a second order Butterworth filter. Baseline activity, estimated as the average activity before stimulus onset (−300 to -50 ms), was subtracted from the pupil diameter time course on a trial-by-trial basis. Next, we z-score normalized the pupil diameter data for each session. Pupil diameter was averaged in 250 ms windows around 500 ms following stimulus onset, 500 ms following cue onset and between 300 to 50 ms before the first-dimming event.

### Analysis of cell type

We distinguished between different cell types based on the duration of the extracellular spike waveform as described in Thiele et al. (2016). Specifically, we classified cells based on the peak-to-trough ratio, i.e. the duration between the peak and the trough of the interpolated (cubic spline) spike waveform. To test whether the distribution of peak-to-trough distance of the spike waveforms was unimodal (null hypothesis) or bimodal, indicating that our distribution contained different cell types, a modified Hartigan’s dip test was used (Ardid et al., 2015; Thiele et al., 2016). We used a cut-off of 250 µs to classify cells as narrow or broad-spiking, as this was where our distribution revealed the main ‘dip’ (Figure 3A-B).

### Fano factor

The variability of neural responses was quantified using Fano factors (*FF*), computed as the ratio between the variance (*σ*^2^) and the mean (*μ*) spike counts within the time window of interest, defined as:

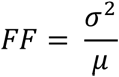

### Drug modulation

The strength of the effect of drug application on neural activity (firing rates) was determined via a drug modulation index (*drugMI*), defined as:

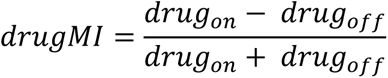

with *drug_on_* as the neural activity when drug was applied, and *drug_off_* the activity when the drug was not applied. This index ranges from -1 to 1, with zero indicating no modulation due to drug application and with positive values indicating higher activity when the drug was applied and conversely, negative values indicating lower activity.

### Quantification of attentional rate modulation

To quantify the difference between neural responses when attention was directed towards the RF versus away from the RF, we computed the area under the receiver operating characteristic (AUROC) curve. Stemming from signal detection theory (Green and Swets, 1966), this measure represents the difference between two distributions as a single scalar value, taking into account both the average difference in magnitude as well as the variability of each distribution. This value indicates how well an ideal observer would be able to distinguish between two distributions, for example the neural response when attention is directed towards versus away from its RF. It is computed by iteratively increasing the threshold and computing the proportion (from the first sample to the threshold) of hits and false alarms (FA), i.e. the correct and false classification as samples belonging to one of the activity distributions. The ROC curve is generated by plotting the proportions of hits against the proportion of FAs, and AUROC is taken as the area under the ROC curve. An AUROC of 0.5 indicates that the two distributions were indistinguishable, whereas an AUROC of 0 or 1 indicates that the two distributions were perfectly separable. As the difference from 0.5 indicates the separability of the distributions, we corrected AUROC values (1-AUROC) for both drug conditions when they were below 0.5 when no drugs were applied, i.e. for those units that displayed higher activity when attention was directed towards the distractors compared to when attention was directed towards the RF.

### Gain variability

Neural activity displays super-Poisson variability (larger variance than the mean), resulting from trial-to-trial changes in excitability, that can be modeled by fitting a negative binomial distribution to the spike rate histogram. This distribution is characterized by a dispersion parameter that captures this additional variability and has been proposed to reflect stimulus-independent modulatory influences on excitability (Goris et al., 2014). For each unit, we fit the distribution of firing rates recorded during the 500 ms before the first dimming with a negative binomial distribution and obtained a gain variance (dispersion) term that captures trial-to-trial changes in excitability, separately for each drug and attention condition (but across stimulus direction conditions).

### Experimental design and statistical analysis

We recorded single (SU, n=40) and multi-unit (MU, n=48) activity (total 88 units; 64 from monkey 1, 24 from monkey 2) from two male rhesus macaque monkeys (*Macaca mulatta,* age 9-11 years, weight 8-12.9 kg). We recorded an additional 12 units during saline-control recordings from one female macaque monkey (11 years, 9.1 kg).

To determine whether DA significantly affected neural activity across the population of units, we used linear mixed-effect models using the R packages *lme4* (Bates et al., 2015) and *lmerTest* (Kuznetsova et al., 2017). The modulation of neural activity (firing rates, Fano Factors or gain variability) was modeled as a linear combination of categorical (effect coded) factors drug (on/off), attention (RF/away), unit type (narrow/broad) and all possible interactions as fixed effects with random intercepts for each unit. We sequentially entered predictors into a hierarchical model and tested the model fit after the addition of each predictor using likelihood ratio tests. For small sample sizes, the *χ*^2^ approximation employed in likelihood ratio tests can lead to misleading conclusions. We therefore additionally report the Kenward-Roger approximation for performing F tests to control for Type I errors (Halekoh and Højsgaard, 2014; Kenward and Roger, 1997; Singmann and Kellen, 2019) using the R package *pbkrtest* (Halekoh and Højsgaard, 2014). To aid interpretation of model fit statistics, we also report Bayes Factors, computed from the sample size, number of predictors and *R*^2^ values (Andraszewicz et al., 2015; Rouder and Morey, 2012) using the R package *BayesFactor* (Morey and Rouder, 2018). Finally, to confirm whether each of the measures had a significant effect on neural activity, we performed “robust regression” based on 5000 bootstrap replicates to calculate the 95% CI around slope estimates for the full model. The reported coefficients are the estimates from the full model and the robust regression. Reported significance values are the results from likelihood ratio tests. We followed these analyses up with linear mixed-effect model tests within each unit type and two-sided paired-sample Wilcoxon signed rank tests.

For comparisons within one recording, e.g. spike rates across trials for different conditions, we used analysis of variance (ANOVA) with three factors: attention (towards/away from the RF), drug (on/off) and stimulus direction. To test whether drug application affected behavioral performance, we used sequential linear mixed effects models with attention and drug as fixed effects and with the recording number as a random effect, to account for the repeated measurements in the data.

To test for significant linear or quadratic trends in the drug dose-response curve, we used sequential linear mixed effects models and likelihood ratio tests. For each drug, we tested whether a first order (linear) polynomial fit was better than a constant (intercept-only) fit and subsequently whether a second order (non-monotonic) polynomial fit was better than a linear fit. The modulation due to drug application of the neural response *y* was modeled as a linear combination of polynomial basis functions of the iontophoretic ejection current *X*:

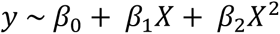

with *β* as the polynomial coefficients. When a significant quadratic relationship was found, we used the two-lines approach to determine whether this relationship was significantly U-shaped (Simonsohn, 2017).

Error bars in all violin plots indicate the interquartile range and the standard error of the mean (SEM) otherwise. We used false discovery rate (FDR) to correct for multiple comparisons.

We selected which cells to include in each of the analyses based on the output of the 3-factor ANOVA described above. For example, if we wanted to investigate whether drug application affected attentional modulation of firing rates, we only included cells that revealed a main or interaction effect for both attention and drug application.

## Data and code availability

Data analyses were performed using custom written scripts in Matlab (the Mathworks) and RStudio (RStudio Team (2016). RStudio: Integrated Development for R. RStudio, Inc., Boston, MA URL http://www.rstudio.com). Violin plots were created using publicly available Matlab code (Bechtold, 2016). Data and analysis scripts necessary to reproduce these results will be made available upon acceptance of this manuscript.

## Acknowledgements

This work was supported by Wellcome trust [093104] (JvK, AT), MRC [MR/P013031/1] (JvK, AT); by a Senior Research Fellowship from the Australian National Health and Medical Research Council (NHMRC) (MAB); and by a strategic research partnership between Newcastle University and Monash University (JvK, MAB, AT).

## Supplementary figures

**Supplementary figure 1.**
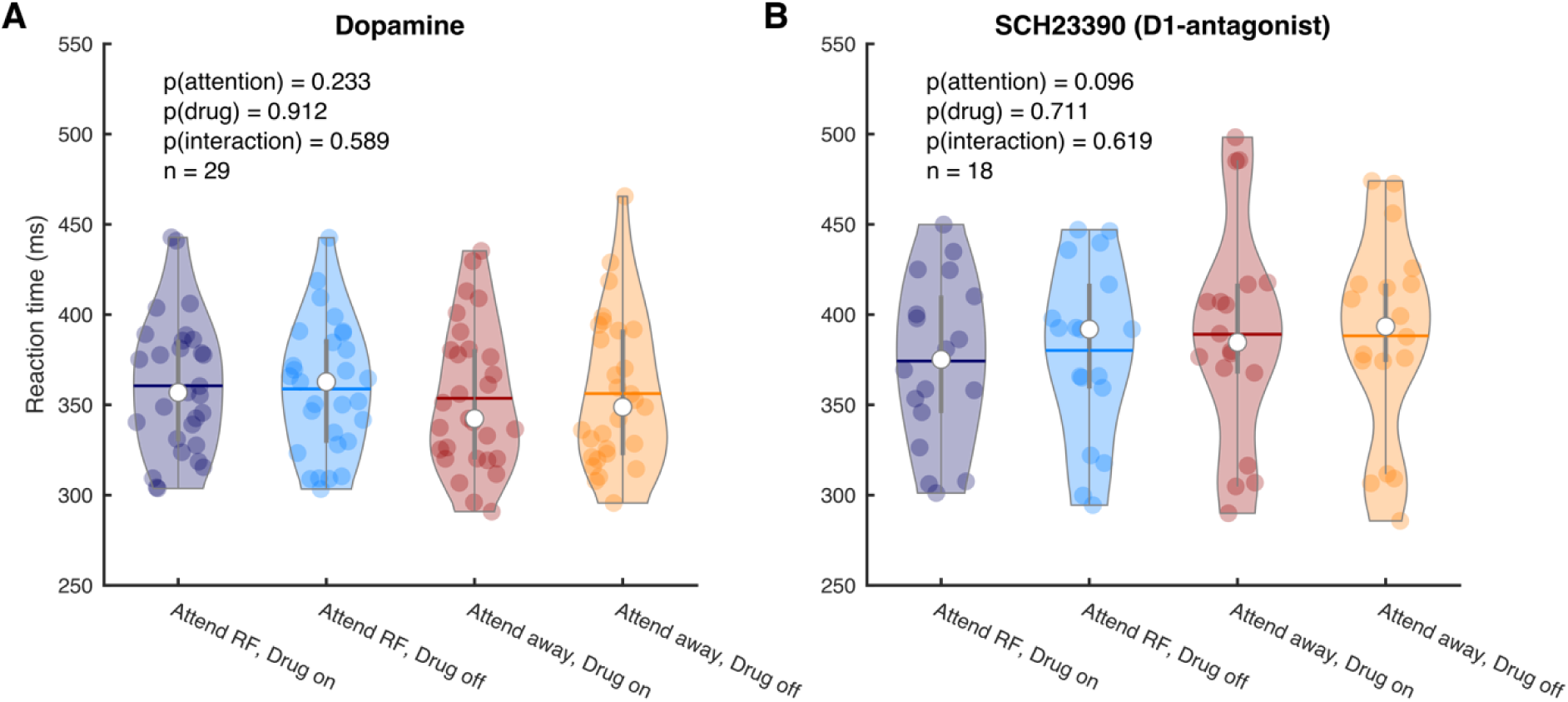
Behavioral performance is unaffected by iontophoretic application of dopaminergic drugs. Average RT on attend RF and attend away trials for the non-specific agonist dopamine (**A**) and the D1R antagonist SCH23390 (**B**). Individual markers represent the average RT during a single recording session. Error bars denote the interquartile range. Horizontal bars denote the mean. Statistics: linear mixed-effects model analysis.

**Supplementary figure 2.**
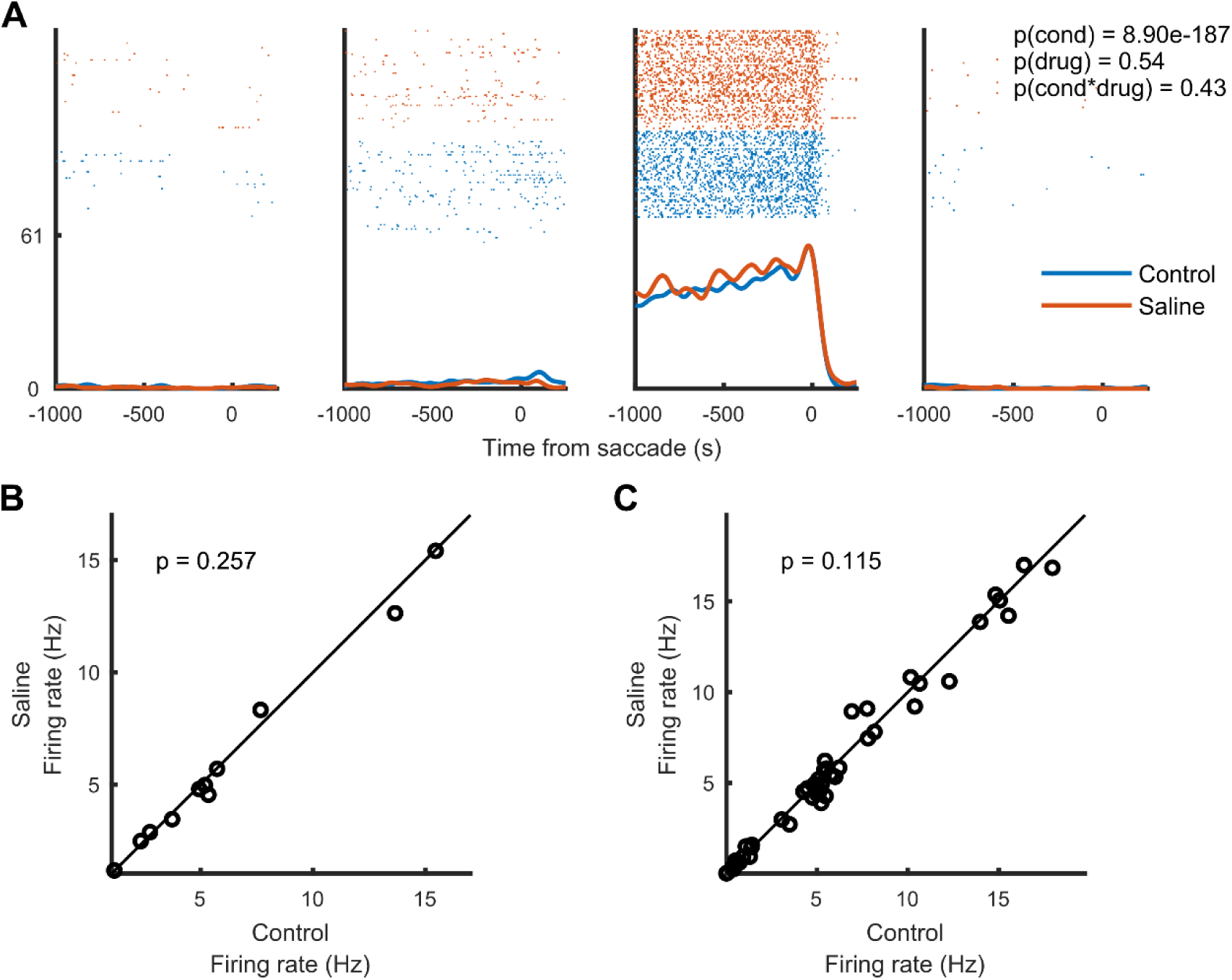
Application of saline with matched pH did not affect firing rates. (**A**) Activity from a representative cell recorded during application of saline (with pH matched to the dopaminergic drugs) whilst the monkey performed a memory-guided saccade task. The four panels correspond to the four quadrants in which the visual stimulus was presented. This cell’s activity, aligned to saccade onset, was significantly modulated by the spatial location of the stimulus/saccade but not by iontophoretic saline application. Statistics: two-factor ANOVA. (**B**) Average firing rates between control and saline conditions. Each marker indicates the average activity of one unit across the four conditions. (**C**) Average firing rates between control and saline conditions. Each marker indicates the average activity of one unit for one of the four conditions. Statistics: two-sided Wilcoxon signed rank test.

**Supplementary figure 3.**
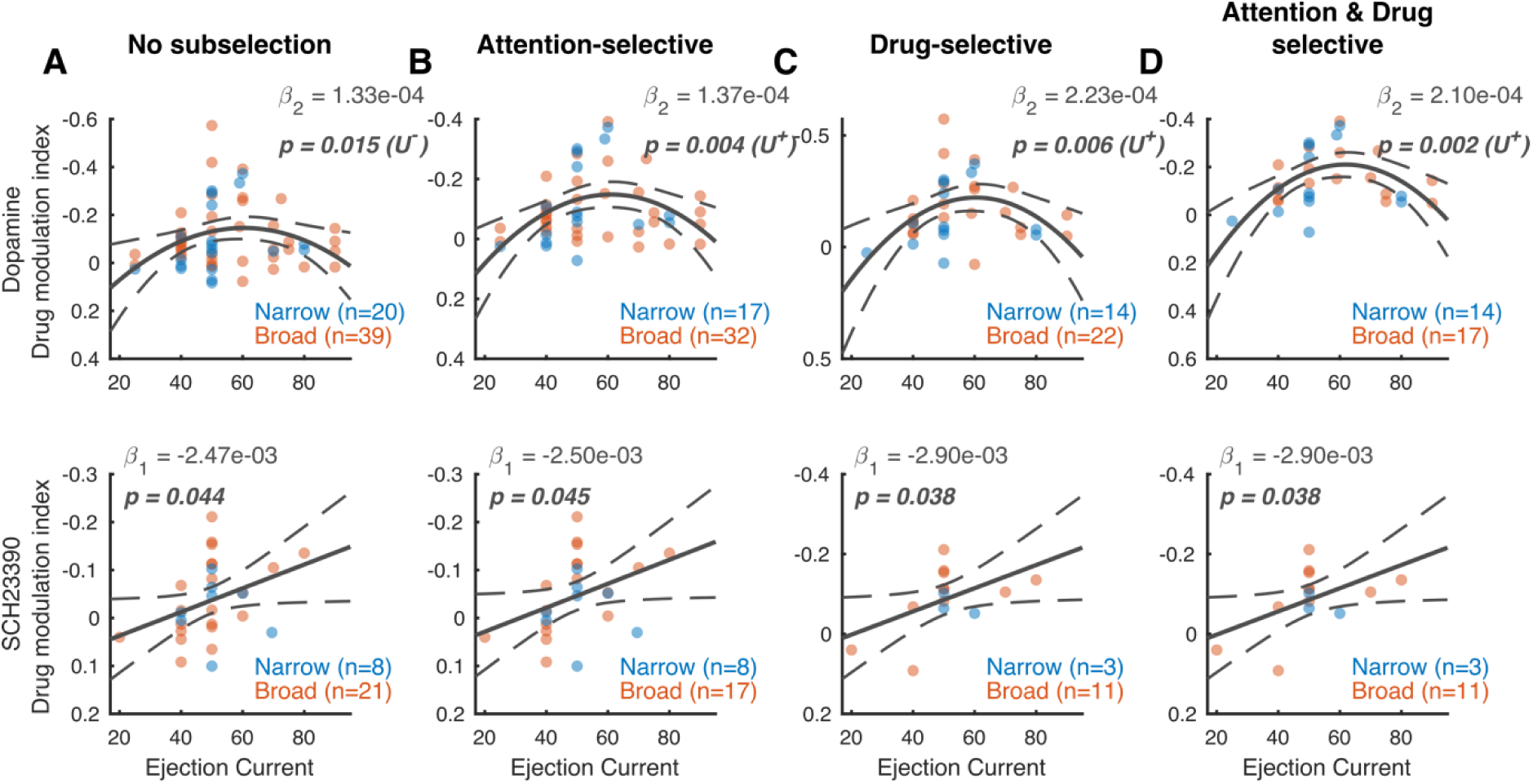
Dose-response curve: drug modulation of firing rates. Drug modulation index plotted against ejection current for the non-specific agonist dopamine (top) and the D1R antagonist SCH23390 (bottom) for (**A**) All units (**B**) units that revealed a main or interaction effect for the factor attention (**C**) units that revealed a main or interaction effect for the factor drug and (**D**) units that revealed a main or interaction effect for the factors attention and drug. Note the reversed y-axis. Solid and dotted lines represent significant model fits (applied to all cells simultaneously) and their 95% confidence intervals, respectively. A monotonic relationship is shown if a first-order fit was better than a constant fit, and a non-monotonic relationship is shown if a second-order fit was better than a linear fit. U^+^ indicates a significant U-shaped relationship. Statistics: linear mixed-effects model analysis. Statistics deemed significant after multiple comparison correction are displayed in italic and boldface fonts.

**Supplementary figure 4.**
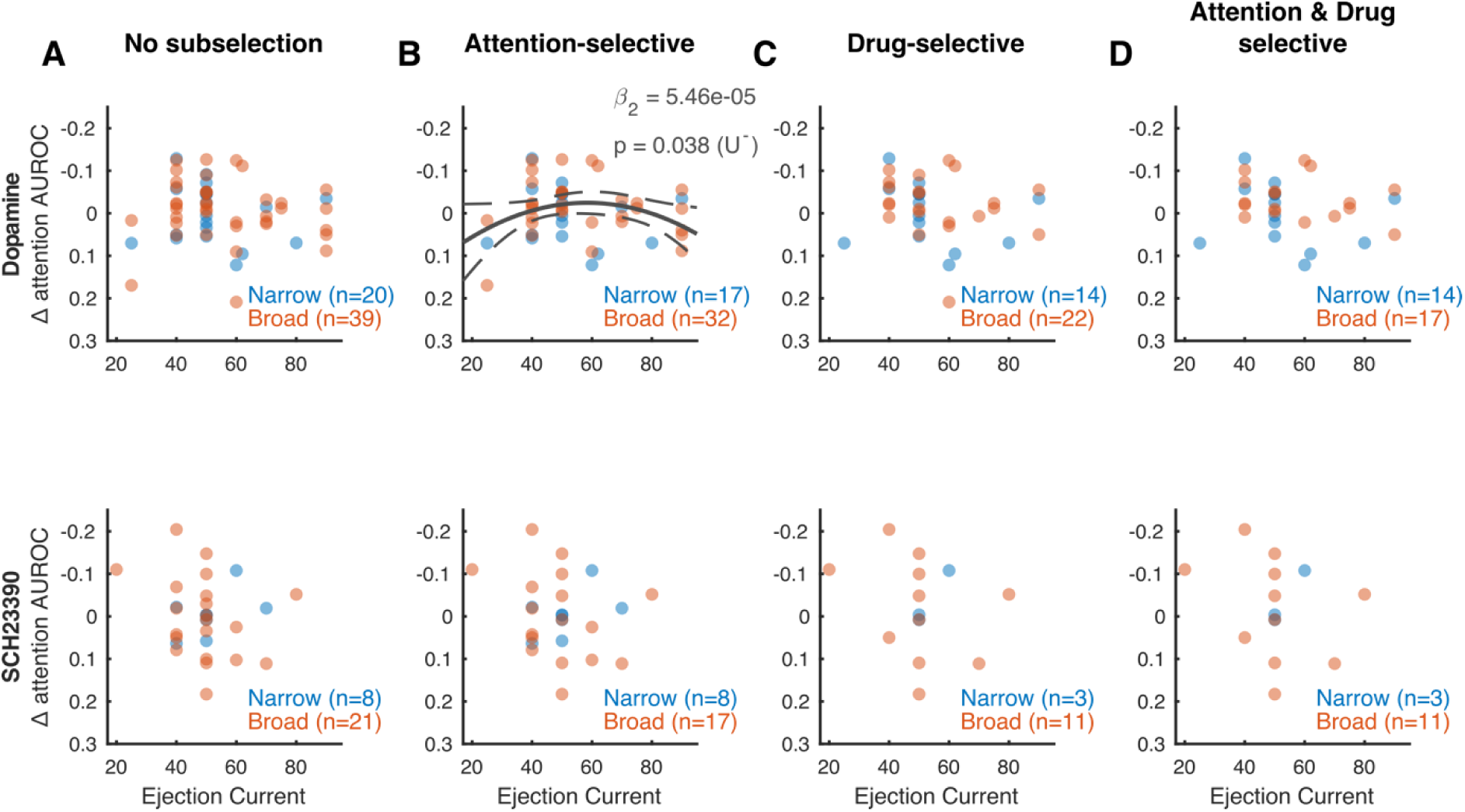
Dose-response curve: drug modulation of attention AUROC values. Attention AUROC difference score (drug-no drug) plotted against ejection current for the non-specific agonist dopamine (top) and the D1R antagonist SCH23390 (bottom) for (**A**) All units (**B**) units that revealed a main or interaction effect for the factor attention (**C**) units that revealed a main or interaction effect for the factor drug and (**D**) units that revealed a main or interaction effect for the factors attention and drug. Note the reversed y-axis. Solid and dotted lines represent significant model fits (applied to all cells simultaneously) and their 95% confidence intervals, respectively. A monotonic relationship is shown if a first-order fit was better than a constant fit, and a non-monotonic relationship is shown if a second-order fit was better than a linear fit. U^+^ indicates a significant U-shaped relationship. Statistics: linear mixed-effects model analysis. Statistics deemed significant after multiple comparison correction are displayed in italic and boldface fonts.

